# A methoxylated flavone from *Artemisia afra* kills *Mycobacterium tuberculosis*

**DOI:** 10.1101/2023.10.11.561885

**Authors:** Joshua J. Kellogg, Maria Natalia Alonso, R. Teal Jordan, Junpei Xiao, Juan Hilario Cafiero, Trevor Bush, Melissa Towler, Pamela Weathers, Scarlet S. Shell

## Abstract

Tuberculosis, caused by Mycobacterium *tuberculosis* (Mtb), is a deadly and debilitating disease globally affecting millions annually. Emerging drug-resistant Mtb strains endanger the efficacy of the current combination therapies employed to treat tuberculosis; therefore, there is an urgent need to develop novel drugs to combat this disease. *Artemisia afra* is used traditionally in southern Africa to treat malaria and recently has shown anti tuberculosis activity. This genus synthesizes a prodigious number of phytochemicals, many of which have demonstrated human health effects. Transcriptomic analysis revealed that *A. afra* exerts different effects on Mtb compared to *A. annua* or the well-known antimalarial artemisinin, suggesting other phytochemicals present in *A. afra* with unique modes of action. A biochemometric study of *A. afra* resulted in the isolation of a methoxylated flavone (**1**), which displayed considerable activity against Mtb strain mc^2^6230. Compound **1** had an MIC of 312.5 μg/mL and yielded no viable colonies after 6 days of treatment. In addition, **1** was effective in killing hypoxic Mtb cultures, with no viable cultures after 2 days of treatment. This suggested that *A. afra* is a source of potentially powerful anti-Mtb phytochemicals with novel mechanisms of action.

## Introduction

Tuberculosis (caused by infections of *Mycobacterium tuberculosis*, Mtb) killed 1.6 million people globally in 2021, and has seen a worrying uptick in incidence and mortality in the last several years, likely due to the SARS-CoV-2 pandemic.^1,2^ To combat this debilitating disease, the World Health Organization (WHO) developed the “End TB Strategy”, which aims to eradicate the disease by 2050.^1^ To meet that goal, a greater emphasis on long-term treatments of tuberculosis is direly needed. Currently, susceptible tuberculosis infections require a regimen consisting of an intensive phase of two months of isoniazid, rifampin, pyrazinamide, and ethambutol followed by a second sustained phase of four months of isoniazid and rifampin combination therapy.^3^ Long periods of multi-drug exposure are needed to slow the emergence of resistance and to kill persistent, non-replicating bacteria, which have greatly reduced drug sensitivity but will cause relapse if not fully cleared. As multidrug resistant (MDR), extensively drug resistant (XDR), and total drug resistant (TDR) strains emerge, current first-line treatments are not sufficient, requiring longer treatment regimens^3,4^ Even in cases of fully drug sensitive TB, the length and complexity of the first-line regimen creates barriers in the resource-limited settings where the disease is most prevalent. To meet the evolving challenge of tuberculosis today and meet the WHO’s goals, newly developed drugs are urgently needed. Drugs that could more rapidly kill non-replicating Mtb would be particularly valuableAfrican wormwood (*A. afra*) is a member of a prolific biosynthetic genus; the Dictionary of Natural Products (http://dnp.chemnetbase.com) contains ca. 1500 records for the genus *Artemisia*. This genus of over 500 species is a source of important bioactive compounds, the most famous of which is artemisinin, the noted antimalarial produced by *Artemisia annua* L.^5–7^ While both *Artemisia* species contain a plethora of specialty molecules, *A. afra* typically contains no detectable artemisinin.^8^ Also similar to *A. annua, A. afra* has a long history of ethnobotanical medicinal use, primarily in southern Africa instead of South Asia.^8^ Intriguingly, recent preliminary reports have suggested that *A. afra* has benefits to clinical TB treatment.^9,10^

In the ever-escalating arms race between pharmaceutical development and infectious agent’s evolution of resistance, natural products have remained at the forefront of antimicrobial discovery.^11,12^ Indeed, a number of natural products from plants have demonstrated efficacy as antimycobacterial agents;^13^ such sustained discovery efforts are crucial for the continued protection of human health against tuberculosis. Dichloromethane (DCM) extracts of *A. annua* and *A. afra* both previously demonstrated bactericidal activity against Mtb.^12,14^ While artemisinin had weak bactericidal activity, it could not fully explain the bactericidal activity of *A. annua*, nor any of the bactericidal activity of *A. afra*, which lacks artemisinin.^14,16^ We therefore hypothesized that there must be some other phytochemicals in *A. afra* that are not artemisinin that are responsible for this bioactivity. *A. afra* retained activity in multiple carbon sources and in hypoxia, a condition frequently encountered by Mtb during human disease.^16^ These observations suggested that *A. afra* may contain compounds with particular clinical potential, as many existing drugs are carbon-source-dependent and/or have poor activity against hypoxia-arrested Mtb.

Here we performed transcriptomic profiling to assess the biological impact of *Artemisia* extracts on Mtb, then investigated the phytochemical content of *A. afra* responsible for its activity. We harnessed biochemometry, fusing untargeted metabolomic profiling with *in vitro* bioassay data to identify potential antimycobacterial compounds. A DCM extract of *A. afra* was fractionated and assayed to determine minimum inhibitory concentration (MIC) as well as bactericidal activity against Mtb. The active fraction (HAB9) was separated chromatographically, and the metabolomic profile and anti-Mtb assay data were combined in a supervised multivariate model, to highlight the presence of a methoxylated flavone structure **1**. The active fraction, containing >97% Compound **1**, was shown to possess bactericidal activity against Mtb both during aerobic growth and in arrest of hypoxia-induced growth.

## Results and Discussion

### The transcriptomic impact of *Artemisia* extracts and artemisinin

Upon exposure to drugs, bacteria generally up- and down-regulate sets of genes as they attempt to cope with the specific stressors imposed by the treatment. Transcriptomic profiling can therefore provide insight into the specific effects of different drugs. To assess and compare the transcriptomic responses of Mtb to *Artemisia* extracts, we performed RNAseq on cultures exposed to DCM extracts of *A. annua* and *A. afra*, as well as to pure artemisinin. Each of the three treatments was applied at three levels: a lethal dose, a sub-lethal dose that inhibited most growth, and a dose that slowed growth but did not stop it. The concentrations of drug and extract that produced these phenotypes were determined by plating for CFU. The transcriptomic impacts of the lethal doses were then assessed after four hours, to prevent confounding by the accumulation of dead cells, while the impacts of inhibitory and sub-inhibitory doses were assessed after 24 hours.

The overall transcriptomic profiles clustered by type of treatment (*A. annua, A. afra*, artemisinin) rather than by the severity of the treatment (Fig 1A). This indicated that the profiles largely reflected treatment-specific responses rather than non-specific responses such as slowed growth that are shared between treatments. The transcriptomic impacts of the three treatments were distinct from one another (Fig 1A), indicating that *A. annua* and *A. afra* affect Mtb through distinct mechanisms, and that *A. annua* has substantial impacts on Mtb through artemisinin-independent mechanisms. The overall transcriptomic impact of artemisinin was smaller than that of the plant extracts, as revealed by clustering (Fig 1A) and by the number of differentially expressed genes compared to no-drug controls (Fig S1 and Table S1). This may reflect artemisinin being a single compound that likely has a single mechanism of action towards aerobically growing cells, while the extracts likely contain multiple compounds with different mechanisms of action.

**Figure 1.**
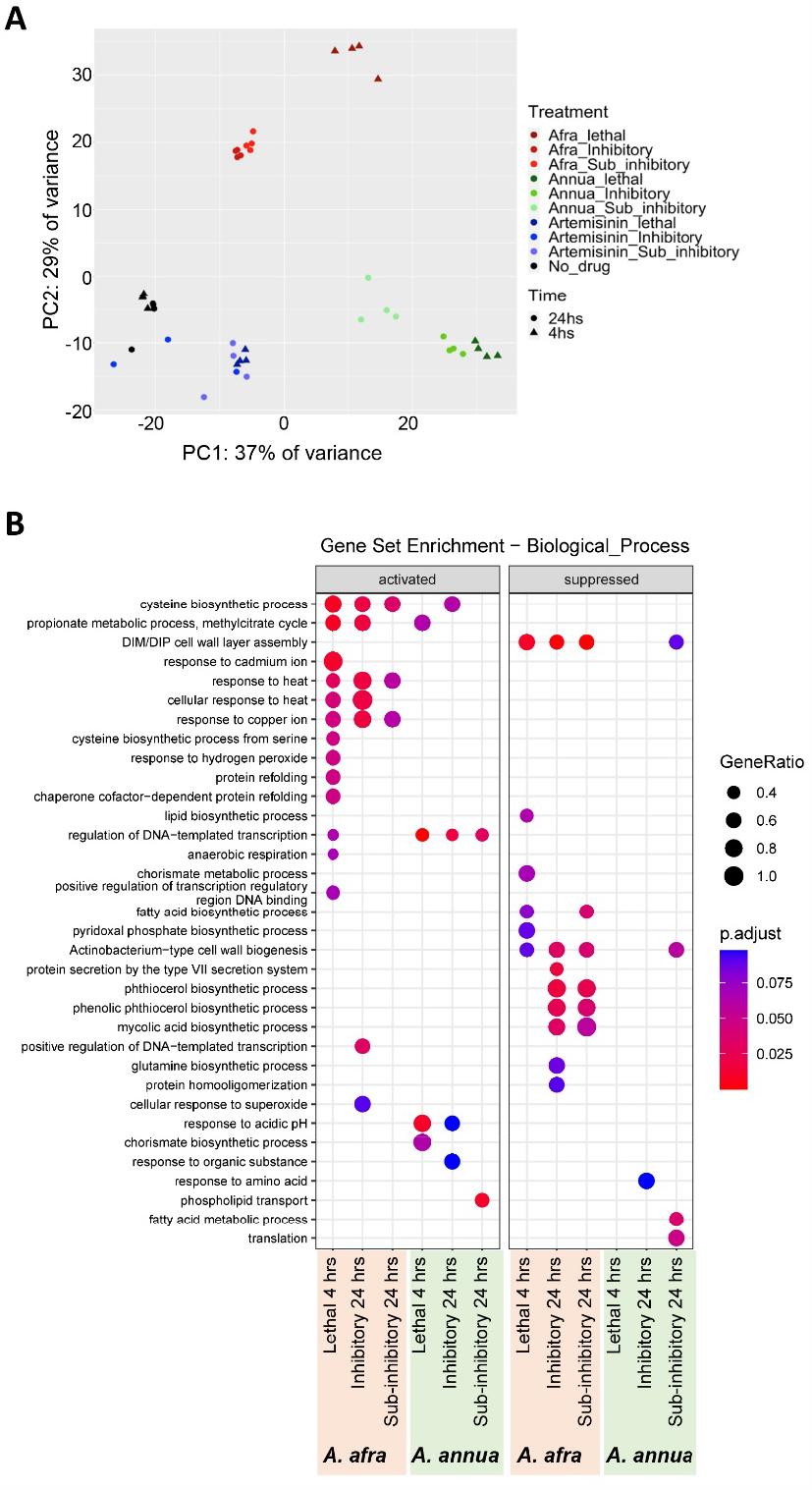
*A. afra, A. annua*, and artemisinin have distinct transcriptomic impacts on Mtb. Aerobically growing Mtb was treated with each extract or compound at lethal doses for four hours or at inhibitory and sub-inhibitory doses for 24 hours. Untreated cultures were harvested at the same two time-points and RNAseq was used to generate transcriptomic profiles. All conditions were tested in quadruplicate. **A**. PCA was done on the read count tables from each sample in each condition, revealing that each treatment clustered separately. **B**. Expression of each gene in each treatment was compared to that in the time-matched control, and differentially expressed genes were subject to Gene Set Enrichment Analysis. Gene sets were GO Biological Process gene lists obtained from AmiGO.^*15*^ “Activated” gene sets had higher expression in the treated samples compared to the controls while “suppressed” genes sets had lower expression. “GeneRatio” indicates the proportion of genes in the set that were differentially expressed in the indicated condition. “p.adjust” is the *P* value of the overrepresentation of genes within the set among the differentially expressed genes, after correction for multiple comparisons.

To investigate the specific physiological impacts of each treatment, we performed Gene Set Enrichment Analysis^17^ to identify pathways that were disproportionately affected. The enriched pathways were largely distinct for *A. annua* and *A. afra* extract treatments, consistent with the idea that their primary mechanisms of action against Mtb are distinct (Fig 1B). No pathways were disproportionately affected by artemisinin. Among the pathways induced by *A. afra* treatment, several were consistent with responses to oxidative stress (eg, cysteine biosynthesis, response to cadmium, response to heat, response to copper, response to hydrogen peroxide, and protein refolding). Notably, many existing antibiotics are thought to induce oxidative stress indirectly, suggesting potential for synergy with *A. afra*. The pathways that were downregulated in response to *A. afra* treatment included several involved in cell envelope biosynthesis. The pathways up- and down-regulated in response to *A. annua* treatment were less easily categorized.

### Fractionation and activity testing of an *A. afra* extract

Given the clearly artemisinin-independent activity of *A. afra*, we sought to identify its active compounds. Flash chromatography was used to create five fractions. One of these, F2, had the lowest MIC against Mtb strain mc^2^6230 (H37Rv ΔRD1 Δ*panCD*)^18^ (Table 1) and also had the most potent bactericidal activity (Fig 2A). However, the fractions were too complex to adequately model their bioactive chemistry via a PLS or OPLS-DA model. Botanicals have been known to possess too many constituents per fraction after a preliminary round of fractionation, so subsequent subfractionation must be undertaken to simplify the chemistry of each sample thereby facilitating a more robust biochemometric model.^19^ Thus, the most active fraction, F2, was selected for further subfractionation on a reverse phase flash system, yielding 16 subfractions (HAB1-10, HA14, 16, 17, and HB13-15). Of these, subfraction HAB9 had among the lowest MICs (Table 1) and strong bactericidal activity against aerobically growing Mtb (Fig 2B). The activity of HAB9 against Mtb in hypoxia-induced growth arrest was so potent that no colonies were observed after two days of treatment, which was the first time-point assessed after addition of the fraction (Fig 2C).

**Table 1.** Minimal inhibitory concentrations (MICs) of *A. afra* extract and fractions.

| Extract, fraction, or compound | MIC<sup>a</sup> (mg/mL; Mtb strain mc<sup>2</sup>6230, aerobic growth) |
| --- | --- |
| *A. afra* crude extract | 1.25-5<sup>b</sup> |
| Fraction F1 | 1.25 |
| **Fraction F2** | 0.3125 |
| Fraction F3 | 1.25 |
| Fraction F4 | >10 |
| Fraction F5 | >10 |
| Subfraction HAB1 | 1.25 |
| Subfraction HAB2 | 1.25 |
| Subfraction HAB3 | 0.625 |
| Subfraction HAB4 | 0.3125 |
| Subfraction HAB5 | 0.3125 |
| Subfraction HAB6 | 0.625 |
| Subfraction HAB7 | 0.3125 |
| Subfraction HAB8 | 0.625 |
| **Subfraction HAB9** | 0.3125 |
| Subfraction HAB10 | 0.3125 |
| Subfraction HA14 | 0.625 |
| Subfraction HA16 | 0.625 |
| Subfraction HA17 | 2.5 |
| Subfraction HB13 | 1.25 |
| Subfraction HB14 | 2.5 |
| Subfraction HB15 | 5 |

<sup>a</sup>MIC was defined as the lowest concentration in a 2-fold dilution series that prevented detectable growth in a resazurin assay.
<sup>b</sup>Results were difficult to interpret due to the intense background coloration of the extract.

**Figure 2.**
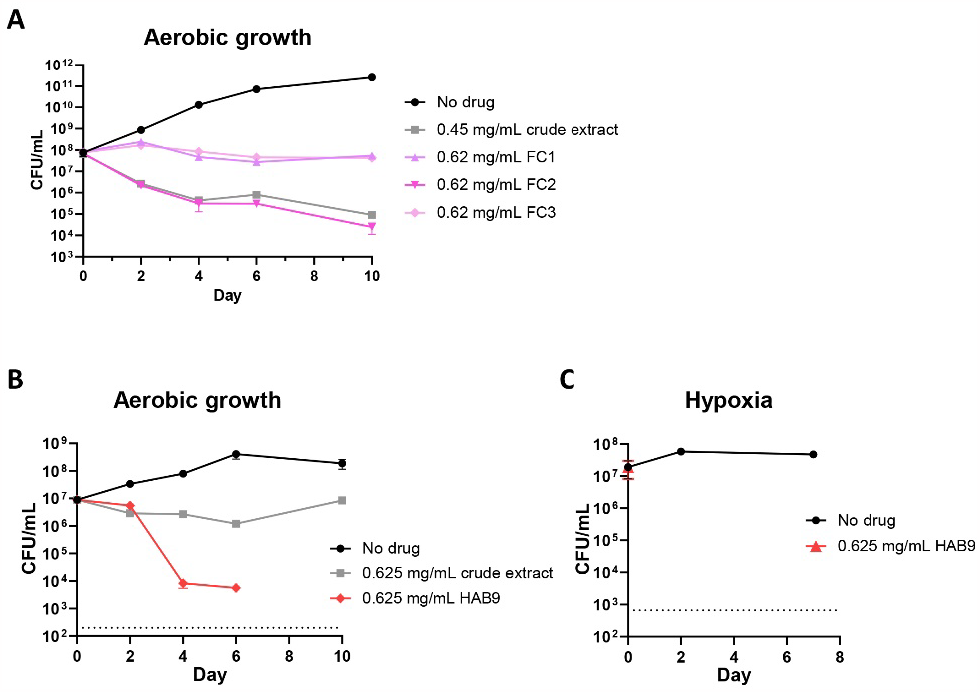
Time-kill curves of Mtb treated with an *A. afra* extract and constituent fractions and subfractions. In all cases, treatment was applied to liquid cultures at Day 0 and aliquots were plated on drug-free solid media at the indicated timepoints. Colonies were enumerated after 3-4 weeks of growth in order to quantify the number of colony forming units (CFU) that had survived drug treatment at each timepoint. All no-drug samples were given DMSO as a vehicle control. **A**. Mtb was exposed to crude DCM extract of *A. afra* and the three flash chromatography fractions with the lowest MICs (see Table 1). **B**. Mtb was exposed to crude DCM extract of *A. afra* and subfraction HAB9. No viable colonies were recovered from cultures treated with HAB9 after six days. The limit of detection is indicated with a dashed line. **C**. Mtb was sealed in vials with 17 mL of culture and 11 mL of headspace. The cultures became hypoxic approximately eight days after sealing as evidenced by methylene blue decoloration. After the cultures had been sealed for 14 days, subfraction HAB9 or the vehicle control DMSO were injected by needle to prevent introduction of oxygen. This was considered day 0 of treatment. No colonies were recovered from cultures treated with HAB9 after two or six days of treatment. The limit of detection is indicated with a dashed line.

### Biochemometric identification of anti-TB compound **1**

Biochemometric modeling of the F2 subfractions was achieved using their untargeted metabolomics profiles of and their classification as ‘active’ or ‘inactive’ based upon the normoxic MIC data. An orthogonal partial least squares-discriminate analysis (OPLS-DA) supervised model led to the correlation and covariance of the features (ions) with the bioactivity categorization. The model gave a good level of fitness (R2X = 0.828, R2Y = 0.778) and good predictivity (Q2 = 0.798), and the resulting S-line plot yielded several metabolites of interest (Figure 3A). The S-line plot highlights variables with a high value of correlation (shown by the red coloration), and covariance (value along the y-axis), play a crucial role in the OPLS-DA discriminatory model between active and inactive. The main covarying feature had an *m/z* of 299.0841, which was observed as the main compound in the subfraction HAB9 (Figure 3B) and was isolated as **1** (Figure 4). Compound **1** comprised 97.4% of HAB9 by mass, and we therefore attribute the activity observed in Figure 2 to Compound **1**.

**Figure 3.**
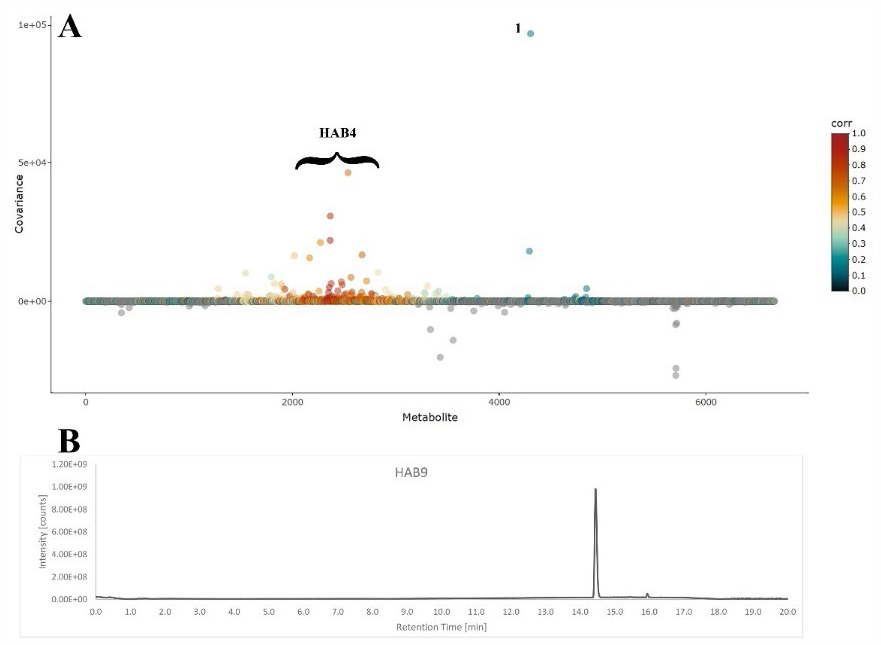
Biochemometric identification of 1. (A) S-line plot illustrating the covariance (y-axis) and correlation (coloration) between features and classification of the OPLS-DA model. Greater values in the covariance and correlation represent features that hold a larger influence in the model generation and discrimination between active and inactive samples. Compound **1** is highlighted, along with ions from another *A. afra* subfraction (HAB4). (B) Extracted ion chromatogram of **1**, *m/z* 299.0841 from subfraction HAB9.

**Figure 4.**
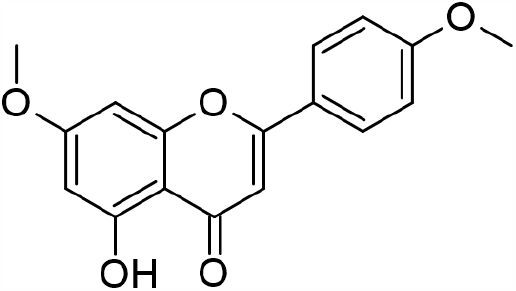
Structure of compound **1**, 5-hydroxy-4’,7-dimethoxyflavone.

Compound **1** had a molecular formula of C_17_H_14_O_5_, based on the protonated HRESIMS ion with an *m/z* of 299.0914. Inspection of the ^1^H, ^13^C, heteronuclear single quantum correlation (HSQC), and heteronuclear multiple bond correlation (HMBC) NMR data permitted the identification of **1** as 5-hydroxy-7-methoxy-2-(4-methoxyphenyl)chromen-4-one (e.g., 5-hydroxy-4’,7-dimethoxyflavone) (See Supplemental Figures S2-S7, Supplemental Table S2). This compound was one of the more prevalent flavonoids in the *A. afra* DCM extract, with a concentration of 2.29 mg/g dried leaf. This flavone was previously demonstrated in other *Artemisia* species,^20–23^ but this is the first reporting of this compound from *A. afra*, and its first reported bioactivity.^24^

### Cytotoxicity studies

Although *A. afra* is not considered a toxic medicinal plant,^25^ the toxicity of HAB9 (**1)**, and other extracts of the plant were evaluated using the whole animal brine shrimp assay (Table 2). The IC_50_ values of the traditional hot water infusion (tea), a DCM extract, and HAB9 were 4.27, >2.5, and 0.023 mg/mL, respectively. The HAB9 fraction, which is identified as 5-hydroxy-4’,7-dimethoxyflavone (**1**), appears to be more toxic in brine shrimp than in mammalian cell cultures where it was deemed nontoxic up to the maximum tested concentration of 0.029 mg/mL (Table 2).^26^ Together these data indicated that toxicity requires further testing.

**Table 2.** Toxicity testing of *Artemisia afra* extracts and fractions in brine shrimp.

| Test system | Sample | IC<sub>50</sub> (mg/mL) | Ref |
| --- | --- | --- | --- |
| Brine shrimp | *A. afra* hot water infusion | 4.27<sup>a</sup> | This study |
|  | *A. afra* DCM extract | >2.50<sup>a,b</sup> |  |
|  | **HAB9** | 0.023 |  |
| In vitro cell lines:<br>HUVECs<br>MCF-7<br>Lu1<br>Col2LNCaP<br>hTERT-RPE1 | 5-hydroxy-4',7-dimethoxyflavone (Compound **1**) | No toxicity within tested range up 0.029 mg/mL | <sup>26</sup> |

<sup>a</sup> Value based on extract of mg dried leaves/mL
<sup>b</sup> Due to solubility issues it was not possible to measure toxicity above this concentration.

## Conclusions

Tuberculosis remains one of the more intractable infectious diseases that plague global society. New drugs are urgently needed to tackle emerging drug resistance, as well as to combat the continual challenge of non-replicating Mtb, a physiological response of Mtb that renders it more resistant to pharmaceutical treatment and makes elimination of the disease more difficult. In our study, *A. afra* extracts showed potent activity against Mtb *in vitro* and were observed to be acting along distinct pathways compared to either *A. annua* or artemisinin. Subsequent biochemometric analysis identified the methoxylated flavone **1** as one of the plant’s phytochemicals active against Mtb. Exposure to **1** substantially reduced viable cultures of Mtb in both replicating and non-replicating stages, which is significant given the insensitivity of non-replicating Mtb to most drugs. While potential toxicity may limit enthusiasm for **1** as a direct anti-Mtb compound, **1** may serve a basis for the design of other novel antimycobacterial drugs. The position of the methoxy groups suggest potential future structure-activity studies to determine optimal positioning and derivatization of flavones and structure modification to improve efficacy and reduce toxicity. Furthermore, there appear to be other fractions containing other *A. afra* phytochemicals that are also antimycobacterial; more work is needed to expand the exploration of *A. afra* for compounds beyond **1** that have potential in the fight against emerging Mtb drug resistance. Collectively, the results presented here provide insight into *A. afra* as a source of effective botanical compounds for the prevention of tuberculosis.

## Experimental Section

### General reagents/materials

All solvents and chemicals used were of reagent or spectroscopic grade, as required, and obtained from VWR (Radnor, PA, USA) or Sigma Aldrich (St. Louis, MO, USA).

### Plant sourcing and identification

For extraction *s*everal kg of dried leaves of *Artemisia afra* Jacq. ex Willd. cv. MAL (voucher FTG181107; B#1RbA.10.12.20) were obtained from Atelier Temenos LLC (Homestead, FL, USA). For transcriptomic, Mtb, and toxicity analyses dried leaves of *A. annua* L. SAM cultivar (voucher MASS 00317314) and the *A. afra* MAL cv. were used. Hot water and DCM extracts were prepared according to Kane et al.^27^ and Martini et al.,^14^ respectively.

### Bacterial strains and culture conditions

*Mycobacterium tuberculosis* (Mtb) strain mc^2^6230 (Δ*panCD*, ΔRD1, like mc^2^6030 but lacking hygromycin resistance^18^) was used for all experiments. Unless indicated otherwise, it was grown in 50 mL conical tubes in Middlebrook 7H9 broth, supplemented with 10% OADC (final concentrations 0.05 g/L oleic acid, 5 g/L bovine serum albumin fraction V, 2 g/L glucose, 0.85 g/L sodium chloride, and 4 mg/L catalase), 0.2% glycerol, 0.05% Tween 80, and 24 μg/mL pantothenate (7H9) at 37°C and 200 rpm, in ambient light. To enumerate CFUs the cultures were plated on Middlebrook 7H10 solid media supplemented with 0.5% glycerol, OADC, and 24 μg/mL pantothenate (7H10).

### Minimal Inhibitory Concentration (MIC) determination

MICs were determined using a resazurin microtiter assay (REMA) as described in Martini et al. (2020).^14^

### Determining the bactericidal activity of *A. afra* and its fractions against Mtb

For aerobic growth experiments, Mtb was grown in 7H9 until the bacteria reached exponential phase. The OD_600_ was adjusted to 0.1 and then *A. afra* plant extract or its fractions were added to triplicate cultures at the indicated concentrations. Triplicate cultures without the addition of drugs were used as a control. Samples were taken at 0, 2, 4, 6, and 10 days of incubation after addition of drugs, then serially diluted in 7H9 and plated on 7H10. Colonies were counted after 21 to 28 days of incubation at 37°C and expressed as CFU/mL culture.

A variation of the Wayne model^28^ was used to determine the bactericidal activity of *A. afra* fraction HAB9 in hypoxic growth conditions. Mtb was grown in 7H9 without oleic acid until bacteria reached exponential phase and then the OD_600_ of the culture was adjusted to 0.1. 17 mL of culture was added to 20 mL serum bottles (Wheaton, actual volume 28 mL). Bottles were sealed using a vial crimper with rubber caps (Wheaton, W224100-181 Stopper, 20 mm) and aluminum caps (Wheaton, 20 mm aluminum seal) and cultures were grown at 37°C with 125 rpm shaking. Methylene blue at 1.5 μg/mL was added to separate cultures and its discoloration was used as an indicator of hypoxic conditions (usually after eight days). Triplicate cultures were opened 14 days after sealing the vials, serially diluted in 7H9, and plated on 7H10 to enumerate the initial CFU/mL. Also 14 days after sealing the vials, *A. afra* subfraction HAB9 diluted in DMSO, or an equivalent volume of DMSO as a no-drug control, was aseptically added to separate cultures at a final concentration 0.625 mg/mL using a syringe and a 30G needle to avoid the introduction of oxygen. After 2 and 7 days of drug treatment, vials were opened and plated to enumerate CFUs as previously stated.

### RNAseq

Mtb cultures were growth to exponential phase, diluted to OD=0.1, and treated for four hours with lethal levels or 24 hours with inhibitory and sub-inhibitory levels of *A. annua* extract, *A. afra* extract, or artemisinin. Untreated cultures were harvested at the same timepoints. Pure artemisinin treatments were 150 μg/mL (lethal, 2X MIC), 100 μg/mL (inhibitory, 1.33X MIC), and 37.5 μg/mL (sub-inhibitory, 0.5X MIC). The extract concentrations for this experiment were expressed in terms of the dry leaf mass used to make the extracts as done in our previous publications,^14,16^ rather than the mass of the extract itself as done for the rest of the experiments in the current study. *A. annua* treatments were the extract from 9 mg dry leaf mass per mL culture (lethal, 2X MIC, and resulting in 75 μg/mL artemisinin), 4.5 mg dry leaf mass per mL culture (inhibitory, MIC, and resulting in 37.5 μg/mL artemisinin), and 2.25 mg dry leaf mass per mL (sub-inhibitory, 0.5X MIC, and resulting in 18.25 μg/mL artemisinin). *A. afra* treatments were the extract from 5 mg dry leaf mass per mL culture (lethal, 2X MIC), 2.5 mg dry leaf mass per mL culture (inhibitory, MIC), and 1.25 mg dry leaf mass per mL (sub-inhibitory, 0.5X MIC). The *A. afra* DCM extract residue weight was approximately 9% of the dry leaf mass, and the actual extract concentrations in the corresponding cultures were therefore approximately 0.45 mg/mL, 0.225 mg/mL, and 0.113 mg/mL, respectively. Quadruplicate cultures were used for each condition. Five mL of culture was used for each 24-hr RNA extraction and 20 mL of culture was used for each 4-hr RNA extraction. RNA was extracted as described^29^ except that 100 μm zirconium beads (OPS Diagnostics) were used for lysis; rRNA was depleted and paired-end Illumina sequencing libraries were constructed as described.^30,31^ Libraries were sequenced at the UMass Medical School Deep Sequencing Core Facility on a HiSeq 4000. Raw data are available on GEO, accession number GSE244235.

Reads were demultiplexed with Cutadapt,^32^ and aligned to the reference genome NC_000962 with BWA mem,^33^ Read count tables were generated with featureCounts^34^ and used for Principal Component Analysis and differential expression analysis with DESeq2.^35^ Each treatment was compared to the time-matched untreated control. Genes with adjusted *p* values <0.05 and log_2_ fold changes >1 or <-1 were considered to be differentially expressed. Gene Set Enrichment Analysis was done with clusterProfiler^17^ using annotations obtained from AmiGO (https://amigo.geneontology.org/amigo).^15^

*Extraction, Fractionation, and Isolation of* ***1***. 2.1 kg *A. afra* leaf material was ground and extracted in dichloromethane (DCM) (1 g:20 mL, VWR International, Radnor, PA, USA) under sonication for 30 minutes. The supernatant was decanted, and the process repeated two additional times. The total supernatant was collected and dried under reduced pressure, yielding 193.5 g crude extract. The crude extract was fractionated on a Biotage Selekt flash chromatography system with a Biotage Sfär HC Duo 350 g column (Biotage, Uppsala, Sweden) and using the ternary solvent system hexane (A), chloroform (B), and methanol (C) and the following (A:B:C) gradient at 200 mL/min: at 0 column volumes (0CV), 100:0:0; 3CV, 0:100:0; 8CV 0:100:0; 9CV 0:95:5; 14CV 0:95:5; 34CV 0:89:20. The flash separation yielded 5 pooled fractions, F1-F5.

F2 was subfractionated on a reverse phase flash column (Biotage Sfär C18 120 g) on the Selekt system with the ternary solvent system acetonitrile (A), water (B), and chloroform (C) and the following (A:B:C) gradient at 50 mL/min: at 0 column volumes (0CV), 2:98:0; 2 CV, 2:98:0; 5 CV, 5:95:0; 25CV, 85:15:0; 30 CV, 85:15:0; 31CV, 100:0:0; 41CV, 75:0:25; 42CV, 60:0:40; 47CV, 0:0:100, 49CV, 0:0:100. Fractionation method was performed twice and overlapping fractions combined, to yield 16 subfractions, HAB1-10, HA14, HA16, HA17, and HB13-15.

*5-Hydroxy-4’,7-dimethoxyflavone (****1***): yellowish solid powder; ^1^H and ^13^C NMR see Supplemental Figures S2-S7 and Supplemental Table S2; HRESIMS *m/z* 299.0914 [M + H]^+^ (calc’d for C_17_H_14_O_5_, 299.0919).

### Toxicity testing

Toxicity of extracts and fractions was measured using the whole organism brine shrimp assay that shows close correlation between plant extracts and rodent data.^36^ To hatch shrimp for testing, Aquatic Foods Great Salt Lake Brine Shrimp Eggs (50 mg) were added to 100 mL of 4% w/v Instant Ocean (Spectrum Brands Pet LLC, AQ-SS1510/1642606) and 3 mg baker’s yeast in a 150 mL beaker, covered in aluminum foil, incubated for 48 hr at 28°C with continuous fluorescent light, and sparged with ambient air at 0.5 LPM. Into individual 3 mL glass vials 10 shrimp were added using a 9 in glass Pasteur pipette and volume brought to 500 μL with Instant Ocean. To each vial a 10 mL aliquot of a 1:2 serial dilution of HAB9 (0-2 mg/mL) diluted in DMSO was added with 490 mL of Instant Ocean to bring the final volume to 1 mL. For hot water and DCM extracts the test range was the extract equivalent of 0-8 mg/mL of dried leaves with dilutions made in water and DMSO, respectively. Hot water extracts were pH adjusted to 7.5 with NaOH. After 24 h incubation, average % viability was calculated by counting the live shrimp per vial. There were three replicates per concentration, and the experiment was repeated three times.

### Mass spectrometry and metabolomics analysis

Ultra-high Pressure (UP) LC-MS data were acquired using an Orbitrap Fusion Lumos Tribrid mass spectrometer (Thermo Scientific, Waltham, MA, USA) with an electrospray ionization source coupled to a Vanquish UHPLC system (Thermo Scientific). Injections (5 μL) were separated by reverse-phase UPLC using an Acquity BEH C18 column (150 × 2.1 mm, 1.7 μm particle size (Waters Corporation, Milford, MA, USA)) held at 55°C with a flow rate of 100 μL/min. The following binary solvent gradient was employed with solvent A (LC-MS grade water with 0.1% formic acid), and solvent B (LC-MS grade acetonitrile): initial isocratic composition of 97:3 (A:B) for 1.0 min, increasing linearly to 85:15 over 4 min, increasing linearly to 5:95 over 11 min, followed by an isocratic hold at 5:95 for 2 min, gradient returned to starting conditions over 0.1 min and held for 1.9 min. The positive ionization mode was utilized over a full scan of m/z 100–1000 with the following settings: spray voltage, 3.5 kV; IT tube temperature, 275°C; vaporizer temperature, 75°C; sheath gas flow and auxiliary gas flow, 25 and 5 units, respectively. Raw spectral data was deposited in the MASSive database (ID: MSV000092962, https://massive.ucsd.edu/ProteoSAFe/dataset.jsp?task=7c2ba865c33c483495d260e2effb4b63).

Data preprocessing was performed with the open-access MZmine3 software.^37^ Steps included peak detection, chromatogram deconvolution, and decomposition, as well as isotope and duplicate peak filtering, followed by sample peak alignment and filtering (Table S3). Peak filtering included removing features not present in all three triplicates of a given sample. Additionally, all features not present in at least one sample at a concentration five-fold higher than the blank were removed from the dataset, and the peak area of the triplicates was averaged. See Supplemental Table S3 for data preprocessing parameters.

### NMR analysis

^1^H and ^13^C NMR spectra were acquired with a Bruker AVIII-500 and Bruker NEO 600 (500 and 600 MHz, respectively; Bruker Corporation, Billerica, MA, USA) equipped with a triple resonance TCI single axis gradient cryoprobe running TopSpin 3.2 operating software. NMR chemical shift values were referenced to residual solvent signals for CDCl_3_ (δ_H_ 7.26 ppm). To collect NMR data, samples were resuspended in CDCl_3_ (Cambridge Isotope Laboratories, Andover, MA, USA), and the following experiments were collected: ^1^H, ^13^C, ^1^H-^1^H COSY, DEPT_135_, HSQC, and HMBC (Figures S1-S6). The NMR data for **1** have been deposited in the Natural Products Magnetic Resonance Database (np-mrd.org).^38^

### Biochemometric analysis

The peak area of extracted ions was used as the variable for chemometrics analysis. SIMCA 17.0 software (Umetrics, Umea, Sweden) was used for chemometrics analysis. Prior to chemometrics analysis, the data were range scaled. No data transformation was performed. Orthogonal partial least squares-discriminate analysis (OPLS-DA) was applied for multivariate modeling. Models were evaluated using the score plot, R^2^X, R^2^Y, and Q^2^ metrics. Cross-validation using the leave-one-out technique was used. An S-line plot was used to identify the important variables responsible for predicting anti-tuberculosis activity in the OPLS-DA model.

### Statistical analysis

For the brine shrimp lethality test, an average of three independent replicate experiments each with three technical replicates per concentration were used to calculate an average IC_50_ using GraphPad Prism version 10.0.0. RNAseq statistical analysis is described in that section.

## Acknowledgements

This work was supported in part by the National Institute of Health’s National Institute for Allergies and Infectious Disease (R21 AI151481) and the USDA National Institute of Food and Agriculture’s Hatch Appropriations (PEN04772). Atelier Temenos provided 2023 summer support for T. Bush. Hadiya Giwa, Tufts University, provided assistance with the brine shrimp assay. The authors would like to thank the Huck Institute’s Metabolomics Core Facility, and Dr. Tapas Mal of the Nuclear Magnetic Resonance Facility at the Pennsylvania State University for supporting instrumentation and data collection. We thank Peter Culviner and Sarah Fortune, Harvard T. H. Chan School of Public Health, for providing the sequences for the rRNA depletion oligos.

## References

(1) World Health Organization. Global Tuberculosis Report 2022; WHO, 2022. https://www.who.int/publications-detail-redirect/9789240061729 (accessed 2023-08-28).

(2) World Health Organization. Tuberculosis deaths rise for the first time in more than a decade due to the COVID-19 pandemic. https://www.who.int/news/item/14-10-2021-tuberculosis-deaths-rise-for-the-first-time-in-more-than-a-decade-due-to-the-covid-19-pandemic (accessed 2023-08-30).

(3) Chauhan, A.; Kumar, M.; Kumar, A.; Kanchan, K. Comprehensive Review on Mechanism of Action, Resistance and Evolution of Antimycobacterial Drugs. Life Sciences 2021, 274, 119301. 10.1016/j.lfs.2021.119301.

(4) Hameed, H. M. A.; Islam, M. M.; Chhotaray, C.; Wang, C.; Liu, Y.; Tan, Y.; Li, X.; Tan, S.; Delorme, V.; Yew, W. W.; Liu, J.; Zhang, T. Molecular Targets Related Drug Resistance Mechanisms in MDR-, XDR-, and TDR-Mycobacterium Tuberculosis Strains. Frontiers in Cellular and Infection Microbiology 2018, 8.

(5) Maciuk, A.; Mazier, D.; Duval, R. Future Antimalarials from Artemisia? A Rationale for Natural Product Mining against Drug-Refractory Plasmodium Stages. Nat. Prod. Rep. 2023, 40 (6), 1130–1144. 10.1039/D3NP00001J.

(6) Trifan, A.; Zengin, G.; Sinan, K. I.; Sieniawska, E.; Sawicki, R.; Maciejewska-Turska, M.; Skalikca-Woźniak, K.; Luca, S. V. Unveiling the Phytochemical Profile and Biological Potential of Five Artemisia Species. Antioxidants 2022, 11 (5), 1017. 10.3390/antiox11051017.

(7) Weathers, P. J. Artemisinin as a Therapeutic vs. Its More Complex Artemisia Source Material. Nat. Prod. Rep. 2022. 10.1039/D2NP00072E.

(8) Liu, N. Q.; Van der Kooy, F.; Verpoorte, R. Artemisia Afra: A Potential Flagship for African Medicinal Plants? South African Journal of Botany 2009, 75 (2), 185–195. 10.1016/j.sajb.2008.11.001.

(9) Gisenya, P.; Ogwang, P. E.; Katotola, E.; Ntezahorigwa, A.; Ndayininahaze, C. Pilot Study to Compare the Use of the National Program Drugs ALONE versus Their Combination with Artemisia Afra Infusions for the Treatment of Pulmonary Tuberculosis. PPIJ 2023, 11 (4), 118–128. 10.15406/ppij.2023.11.00410.

(10) Daddy, B.; Lutgen, P.; Gisenya, P. Breakthrough against Tuberculosis: High Efficacy of Artemisia Afra Infusions. Pharm Pharmacol Int J 2021, Volume 9 (Issue 2). 10.15406/ppij.2021.09.00328.

(11) Newman, D. J.; Cragg, G. M. Natural Products as Sources of New Drugs over the Nearly Four Decades from 01/1981 to 09/2019. Journal of Natural Products 2020, 83 (3), 770–803. 10.1021/acs.jnatprod.9b01285.

(12) Woo, S.; Marquez, L.; Crandall, W. J.; Risener, C. J.; Quave, C. L. Recent Advances in the Discovery of Plant-Derived Antimicrobial Natural Products to Combat Antimicrobial Resistant Pathogens: Insights from 2018– 2022. Nat. Prod. Rep. 2023, 40 (7), 1271–1290. 10.1039/D2NP00090C.

(13) Santhosh, R. S.; Suriyanarayanan, B. Plants: A Source for New Antimycobacterial Drugs. Planta Med 2014, 80 (1), 9–21. 10.1055/s-0033-1350978.

(14) Martini, M. C.; Zhang, T.; Williams, J. T.; Abramovitch, R. B.; Weathers, P. J.; Shell, S. S. Artemisia Annua and Artemisia Afra Extracts Exhibit Strong Bactericidal Activity against Mycobacterium Tuberculosis. Journal of Ethnopharmacology 2020, 262, 113191. 10.1016/j.jep.2020.113191.

(15) Carbon, S.; Ireland, A.; Mungall, C. J.; Shu, S.; Marshall, B.; Lewis, S.; the AmiGO Hub; the Web Presence Working Group. AmiGO: Online Access to Ontology and Annotation Data. Bioinformatics 2009, 25 (2), 288–289. 10.1093/bioinformatics/btn615.

(16) Kiani, B. H.; Alonso, M. N.; Weathers, P. J.; Shell, S. S. Artemisia Afra and Artemisia Annua Extracts Have Bactericidal Activity against Mycobacterium Tuberculosis in Physiologically Relevant Carbon Sources and Hypoxia. Pathogens 2023, 12 (2), 227. 10.3390/pathogens12020227.

(17) Wu, T.; Hu, E.; Xu, S.; Chen, M.; Guo, P.; Dai, Z.; Feng, T.; Zhou, L.; Tang, W.; Zhan, L.; Fu, X.; Liu, S.; Bo, X.; Yu, G. clusterProfiler 4.0: A Universal Enrichment Tool for Interpreting Omics Data. Innovation (Camb) 2021, 2 (3), 100141. 10.1016/j.xinn.2021.100141.

(18) Sambandamurthy, V. K.; Derrick, S. C.; Hsu, T.; Chen, B.; Larsen, M. H.; Jalapathy, K. V.; Chen, M.; Kim, J.; Porcelli, S. A.; Chan, J.; Morris, S. L.; Jacobs, W. R. Mycobacterium Tuberculosis ΔRD1 ΔpanCD: A Safe and Limited Replicating Mutant Strain That Protects Immunocompetent and Immunocompromised Mice against Experimental Tuberculosis. Vaccine 2006, 24 (37), 6309–6320. 10.1016/j.vaccine.2006.05.097.

(19) Britton, E. R.; Kellogg, J. J.; Kvalheim, O. M.; Cech, N. B. Biochemometrics to Identify Synergists and Additives from Botanical Medicines: A Case Study with Hydrastis Canadensis (Goldenseal). Journal of Natural Products 2018, 81 (3), 484–493. 10.1021/acs.jnatprod.7b00654.

(20) Talzhanov, N. A.; Sadyrbekov, D. T.; Smagulova, F. M.; Mukanov, R. M.; Raldugin, V. A.; Shakirov, M. M.; Tkachev, A. V.; Atazhanova, G. A.; Tuleuov, B. I.; Adekenov, S. M. Components of Artemisia Pontica. Chem Nat Compd 2005, 41 (2), 178–181. 10.1007/s10600-005-0107-x.

(21) Valant-Vetschera, K. M.; Wollenweber, E. Flavonoid Aglycones from the Leaf Surfaces of Some Artemisia Spp. (Compositae-Anthemideae). Zeitschrift für Naturforschung C 1995, 50 (5–6), 353–357. 10.1515/znc-1995-5-604.

(22) Valant-Vetschera, K. M.; Fischer, R.; Wollenweber, E. Exudate Flavonoids in Species of Artemisia (Asteraceae—Anthemideae): New Results and Chemosystematic Interpretation. Biochemical Systematics and Ecology 2003, 31 (5), 487–498. 10.1016/S0305-1978(02)00178-3.

(23) Chumbalov, T. K.; Fadeeva, O. V. Flavonoids ofArtemisia Transiliensis. Chem Nat Compd 1969, 5 (5), 364–364. 10.1007/BF00595086.

(24) PubChem. Apigenin 7,4’-dimethyl ether. https://pubchem.ncbi.nlm.nih.gov/compound/5281601 (accessed 2023-08-30).

(25) Kane, N. F.; Kyama, M. C.; Nganga, J. K.; Hassanali, A.; Diallo, M.; Kimani, F. T. Acute Toxicity Effect of Artemisia Afra Plant Extracts on the Liver, Kidney, Spleen and in Vivo Antimalarial Assay on Swiss Albino Mice. Advances in Bioscience and Bioengineering 2019, 7 (4), 64. 10.11648/j.abb.20190704.12.

(26) Jones, W. P.; Lobo-Echeverri, T.; Mi, Q.; Chai, H.-B.; Soejarto, D. D.; Cordell, G. A.; Swanson, S. M.; Kinghorn, A. D. Cytotoxic Constituents from the Fruiting Branches of Callicarpa Americana Collected in Southern Florida,1. J. Nat. Prod. 2007, 70 (3), 372–377. 10.1021/np060534z.

(27) Kane, N. F.; Kiani, B. H.; Desrosiers, M. R.; Towler, M. J.; Weathers, P. J. Artemisia Extracts Differ from Artemisinin Effects on Human Hepatic CYP450s 2B6 and 3A4 in Vitro. Journal of Ethnopharmacology 2022, 298, 115587. 10.1016/j.jep.2022.115587.

(28) Wayne, L. G.; Hayes, L. G. An in Vitro Model for Sequential Study of Shiftdown of Mycobacterium Tuberculosis through Two Stages of Nonreplicating Persistence. Infect Immun 1996, 64 (6), 2062–2069. 10.1128/iai.64.6.2062-2069.1996.

(29) Bar-Oz, M.; Martini, M. C.; Alonso, M. N.; Meir, M.; Lore, N. I.; Miotto, P.; Riva, C.; Angala, S. K.; Xiao, J.; Masiello, C. S.; Misiakou, M.-A.; Sun, H.; Moy, J. K.; Jackson, M.; Johansen, H. K.; Cirillo, D. M.; Shell, S. S.; Barkan, D. The Small Non-Coding RNA B11 Regulates Multiple Facets of Mycobacterium Abscessus Virulence. PLOS Pathogens 2023, 19 (8), e1011575. 10.1371/journal.ppat.1011575.

(30) Culviner, P. H.; Laub, M. T. Global Analysis of the E. Coli Toxin MazF Reveals Widespread Cleavage of mRNA and the Inhibition of rRNA Maturation and Ribosome Biogenesis. Molecular Cell 2018, 70 (5), 868–880.e10. 10.1016/j.molcel.2018.04.026.

(31) Culviner, P. H.; Guegler, C. K.; Laub, M. T. A Simple, Cost-Effective, and Robust Method for rRNA Depletion in RNA-Sequencing Studies. mBio 2020, 11 (2), 10.1128/mbio.00010-20. 10.1128/mbio.00010-20.

(32) Martin, M. Cutadapt Removes Adapter Sequences from High-Throughput Sequencing Reads. EMBnet.journal 2011, 17 (1), 10–12. 10.14806/ej.17.1.200.

(33) Li, H.; Durbin, R. Fast and Accurate Long-Read Alignment with Burrows-Wheeler Transform. Bioinformatics 2010, 26 (5). 10.1093/bioinformatics/btp698.

(34) Liao, Y.; Smyth, G. K.; Shi, W. featureCounts: An Efficient General Purpose Program for Assigning Sequence Reads to Genomic Features. Bioinformatics 2014, 30 (7), 923–930. 10.1093/bioinformatics/btt656.

(35) Love, M. I.; Huber, W.; Anders, S. Moderated Estimation of Fold Change and Dispersion for RNA-Seq Data with DESeq2. Genome Biology 2014, 15 (12), 550. 10.1186/s13059-014-0550-8.

(36) Lagarto Parra, A.; Silva Yhebra, R.; Guerra Sardiñas, I.; Iglesias Buela, L. Comparative Study of the Assay of Artemia Salina L. and the Estimate of the Medium Lethal Dose (LD50 Value) in Mice, to Determine Oral Acute Toxicity of Plant Extracts. Phytomedicine 2001, 8 (5), 395–400. 10.1078/0944-7113-00044.

(37) Schmid, R.; Heuckeroth, S.; Korf, A.; Smirnov, A.; Myers, O.; Dyrlund, T. S.; Bushuiev, R.; Murray, K. J.; Hoffmann, N.; Lu, M.; Sarvepalli, A.; Zhang, Z.; Fleischauer, M.; Dührkop, K.; Wesner, M.; Hoogstra, S. J.; Rudt, E.; Mokshyna, O.; Brungs, C.; Ponomarov, K.; Mutabdžija, L.; Damiani, T.; Pudney, C. J.; Earll, M.; Helmer, P. O.; Fallon, T. R.; Schulze, T.; Rivas-Ubach, A.; Bilbao, A.; Richter, H.; Nothias, L.-F.; Wang, M.; Orešič, M.; Weng, J.-K.; Böcker, S.; Jeibmann, A.; Hayen, H.; Karst, U.; Dorrestein, P. C.; Petras, D.; Du, X.; Pluskal, T. Integrative Analysis of Multimodal Mass Spectrometry Data in MZmine 3. Nat Biotechnol 2023, 41 (4), 447–449. 10.1038/s41587-023-01690-2.

(38) Wishart, D. S.; Sayeeda, Z.; Budinski, Z.; Guo, A.; Lee, B. L.; Berjanskii, M.; Rout, M.; Peters, H.; Dizon, R.; Mah, R.; Torres-Calzada, C.; Hiebert-Giesbrecht, M.; Varshavi, D.; Varshavi, D.; Oler, E.; Allen, D.; Cao, X.; Gautam, V.; Maras, A.; Poynton, E. F.; Tavangar, P.; Yang, V.; van Santen, J. A.; Ghosh, R.; Sarma, S.; Knutson, E.; Sullivan, V.; Jystad, A. M.; Renslow, R.; Sumner, L. W.; Linington, R. G.; Cort, J. R. NP-MRD: The Natural Products Magnetic Resonance Database. Nucleic Acids Research 2022, 50 (D1), D665–D677. 10.1093/nar/gkab1052.

