## Supplemental Information for "A methoxylated flavone from *Artemisia afra* kills *Mycobacterium tuberculosis*"

**Figure S1. Upset plot summarizing the commonalities in gene expression changes across treatments.** Genes were classified as differentially expressed if their log2 fold change in mRNA abundance in a treatment compared to its control was >1 or <-1 with an adjusted p value < 0.05. The total number of differentially expressed genes in each treatment is shown on the lower left ("set size"). "Inclusive intersection size" (main histogram) indicates the number of genes that were differentially expressed in any two or three treatments. "Inclusive" means that genes were included in a set even if they also were present in other sets. The plot was made with *ComplexUpset* R package version 0.5 18.

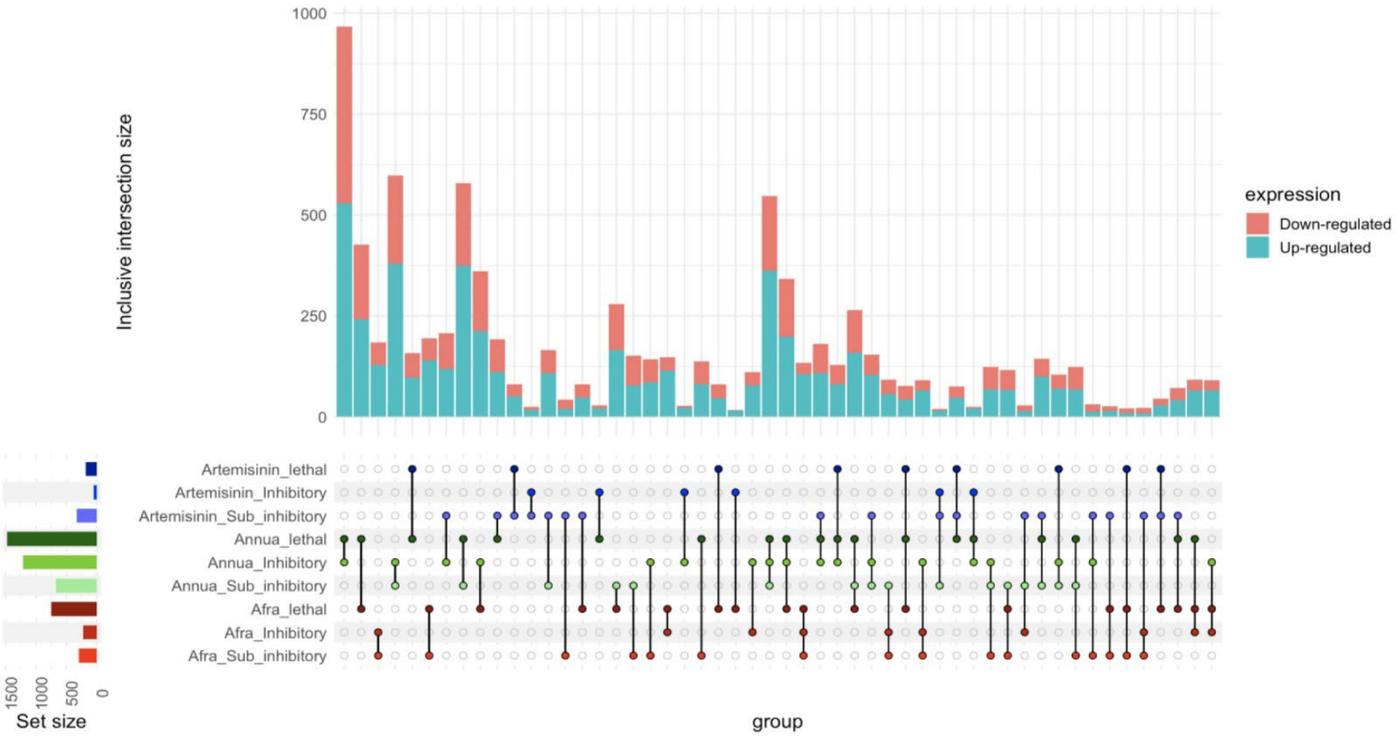


**Figure S2. 1H NMR spectrum for compound 1.**

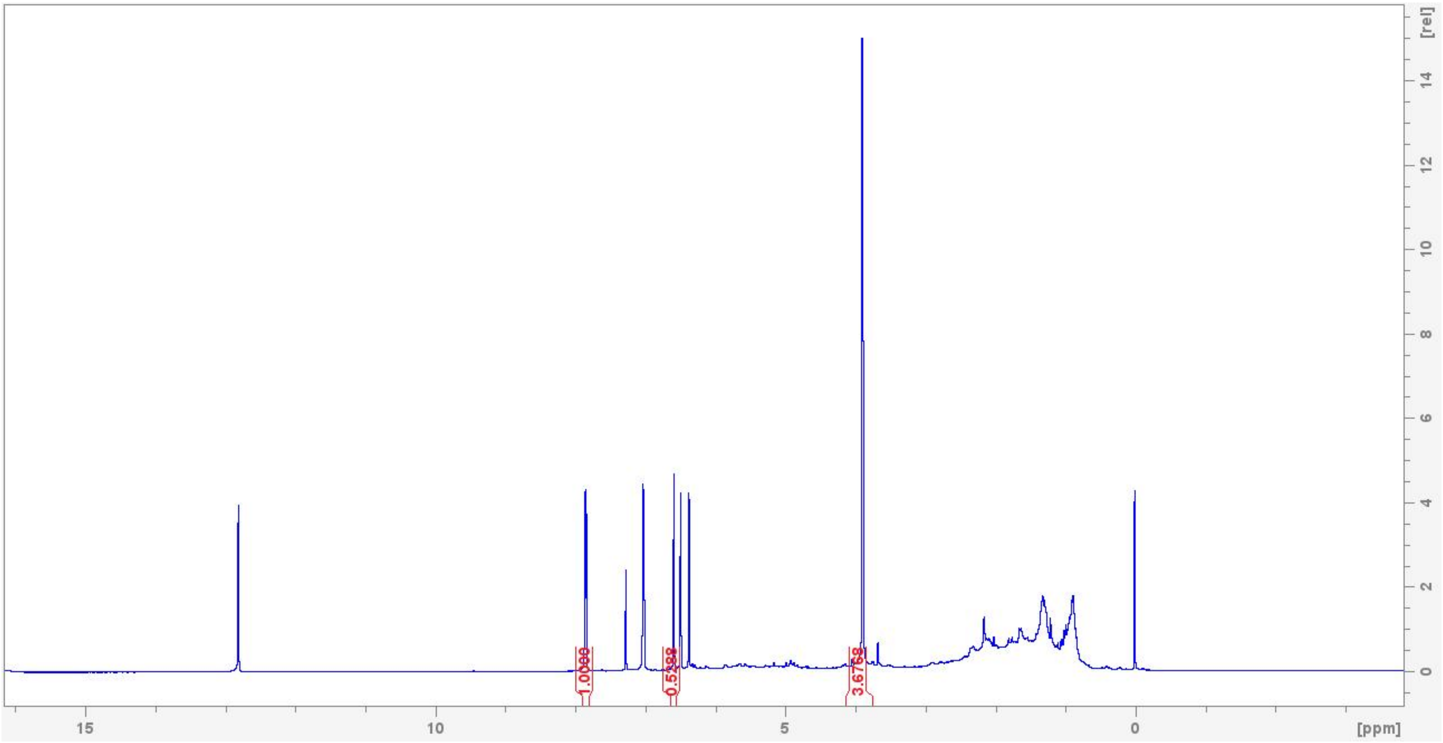


**Figure S3. 13C NMR spectrum for compound 1.**

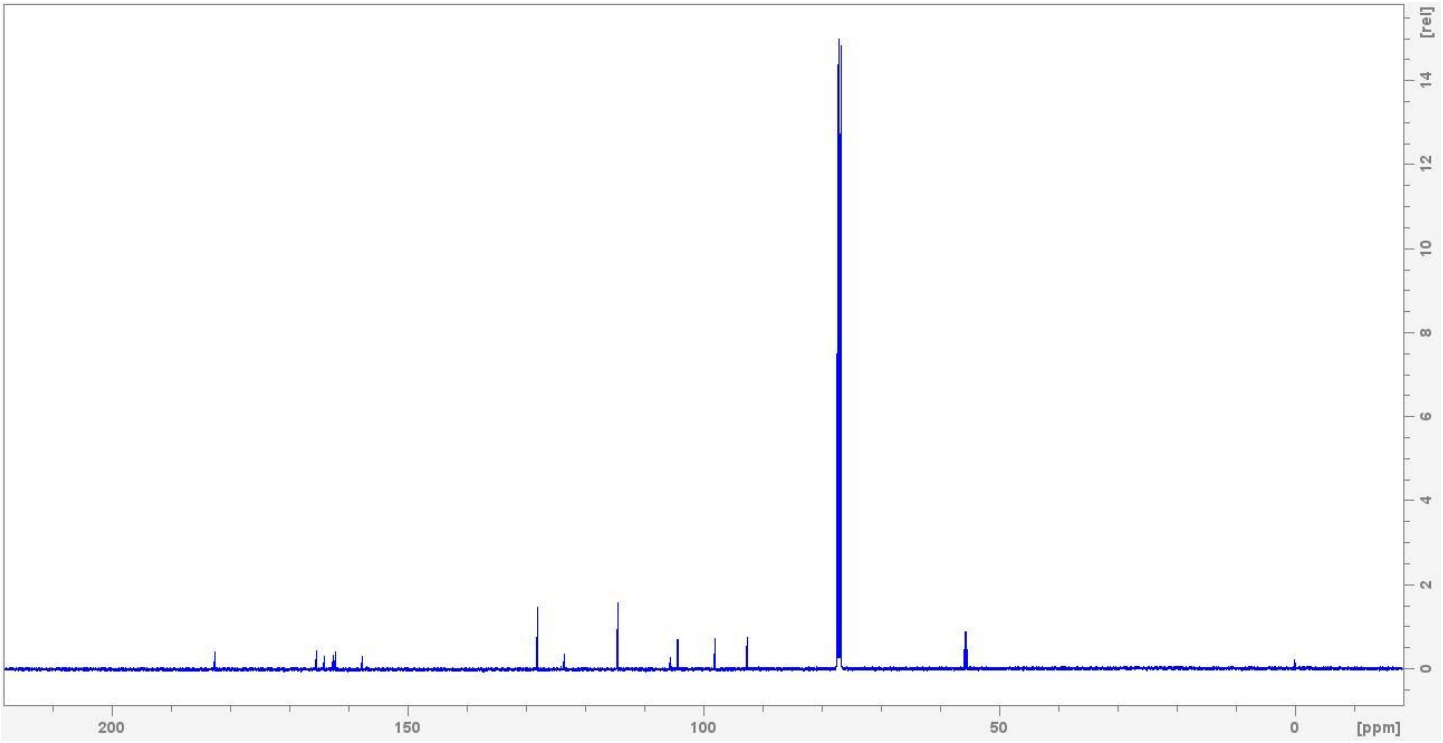

**Figure S4. DEPT-135 spectrum for compound 1.**

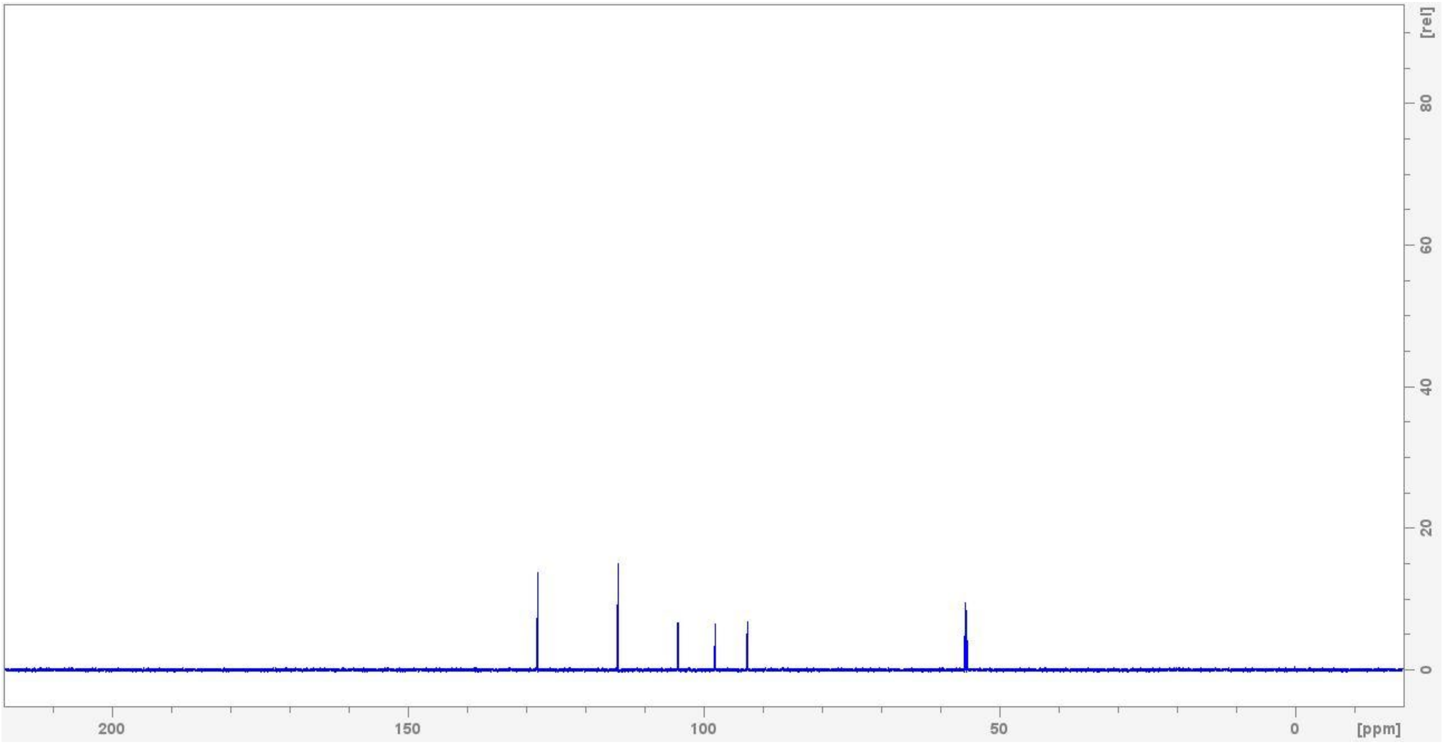


**Figure S5. COSY spectrum for compound 1.**

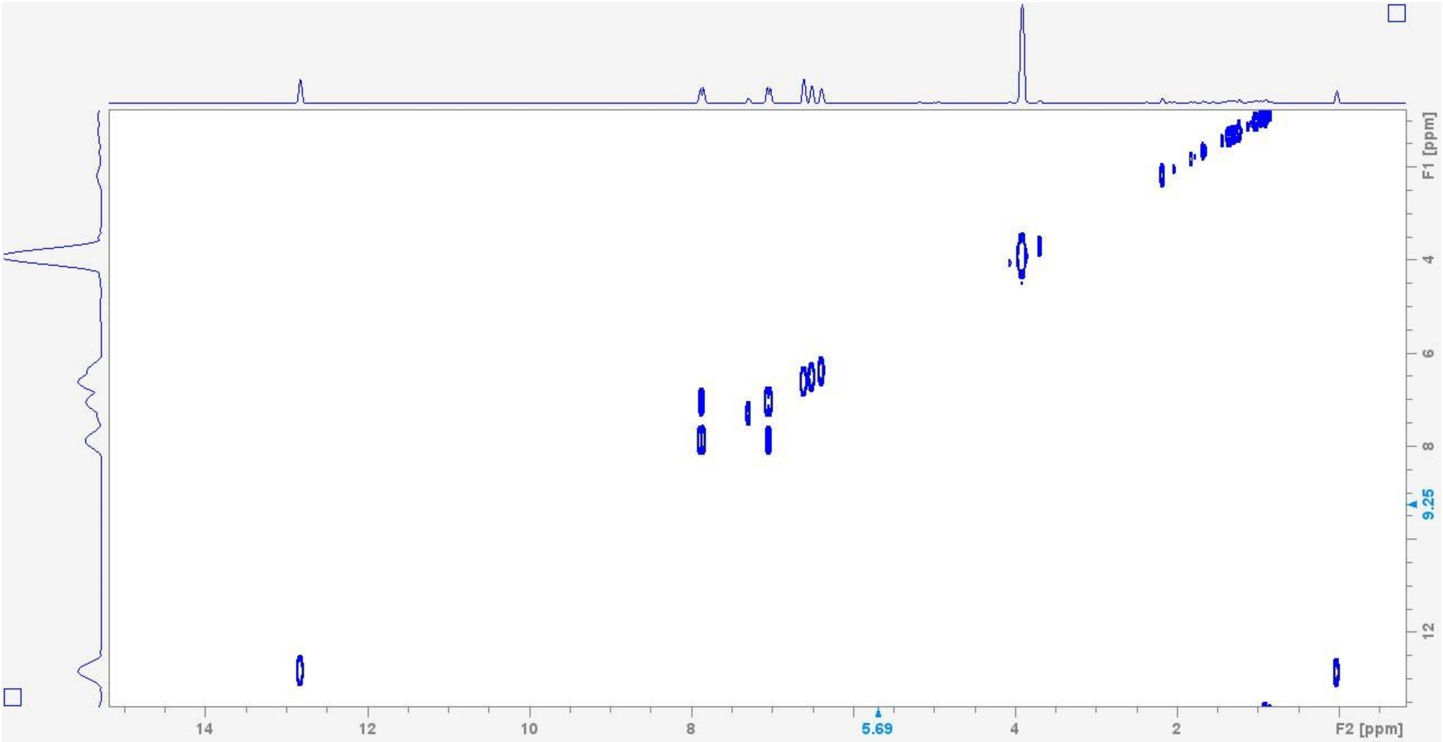

**Figure S6. HSQC spectrum for compound 1.**

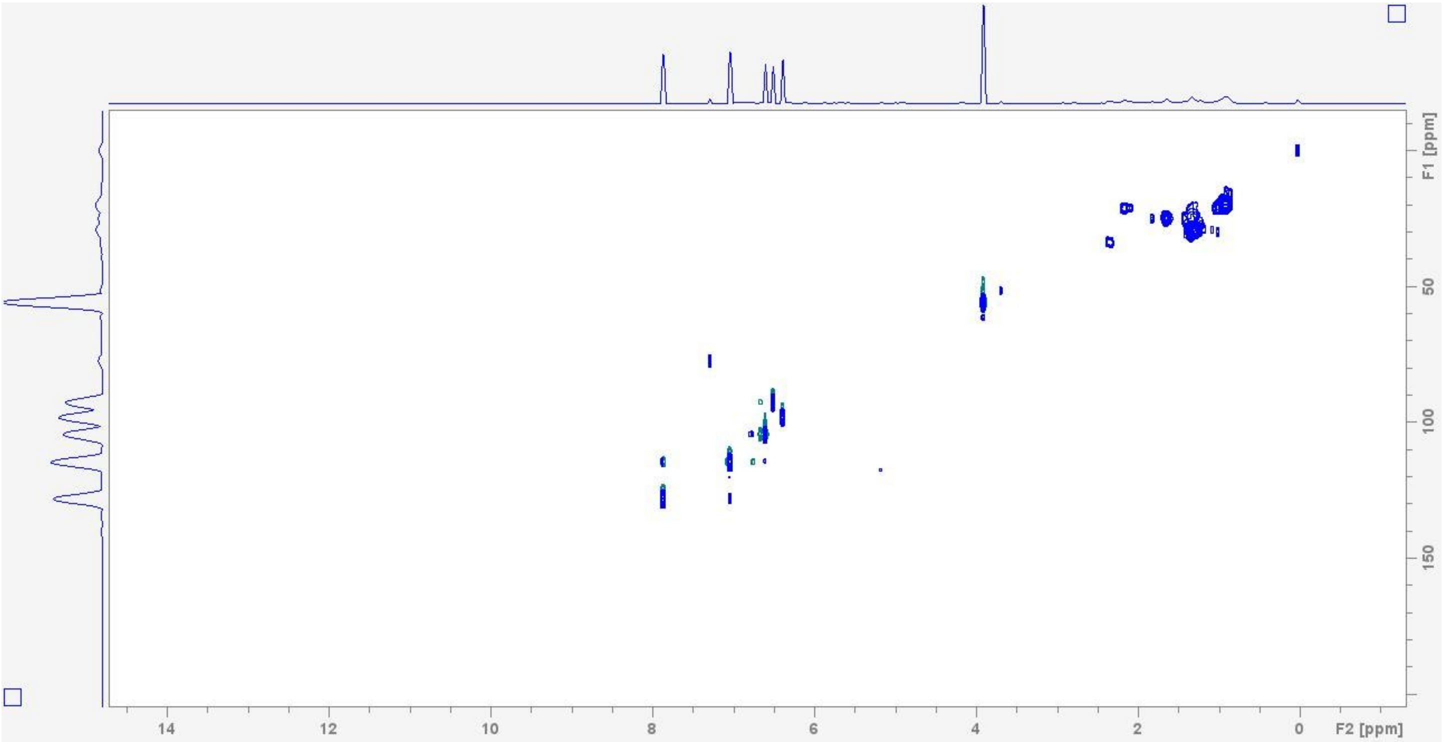


**Figure S7. HMBC spectrum for compound 1.**

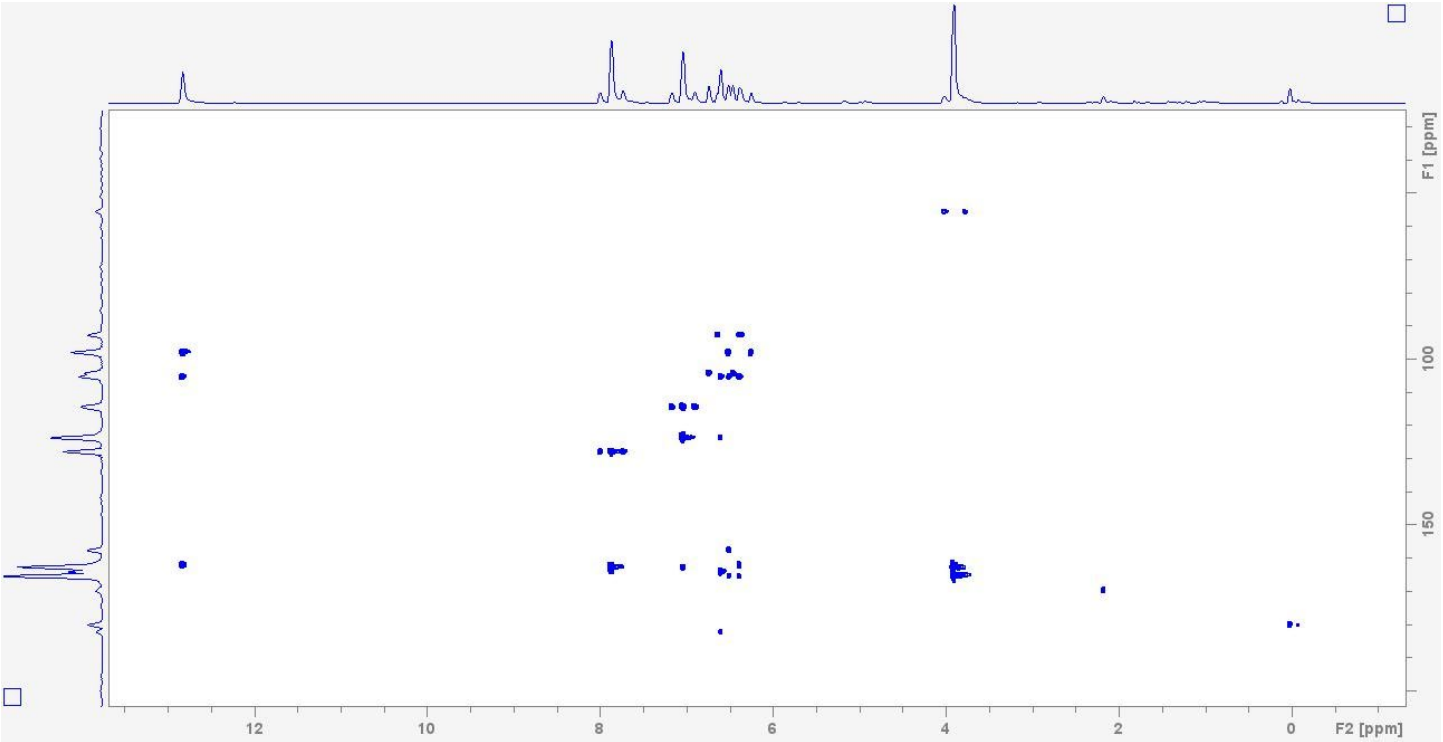

**Table S2. NMR data for compound 1.**

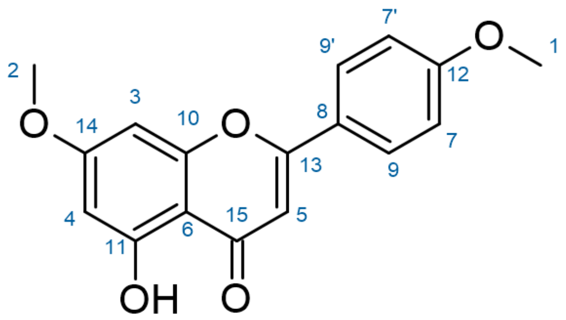


| **Position** | $\delta_C$ | **Type** | $\delta_H$ (J in Hz) | **COSY correlation** | **HMBC correlation** |
|---|---|---|---|---|---|
| **1** | 55.67 | CH3 | 3.88, s | | 12 |
| **2** | 55.93 | CH3 | 3.89, s | | 14 |
| **3** | 92.76 | CH | 6.48, q (2) | | 3,6,10,14 |
| **4** | 98.19 | CH | 6.36, q (2) | | 6,11,14 |
| **5** | 104.47 | CH | 6.58, m | | 6,8,13,15 |
| **6** | 105.68 | C | | | |
| **7, 7'** | 114.64 | CH2 | 7.03, d (8) | 9 | 7'/7,8,12 |
| **8** | 123.69 | C | | | |
| **9,9'** | 128.19 | CH2 | 7.86, d (8) | 7 | 9'/9,12 |
| **10** | 157.84 | C | | | |
| **11** | 162.30 | C | | | |
| **12** | 162.74 | C | | | |
| **13** | 164.18 | C | | | |
| **14** | 165.58 | C | | | |
| **15** | 182.60 | C | | | |
| **OH** | -- | OH | 12.80, s | | 4,6,11 |

**Table S3. MZmine parameters for untargeted metabolomics processing**

| | |
|---|---|
| **MZmine version** | 3.4.27 |
| **Mass Detection** | |
| a. Mass detector | Centroid |
| b. Noise level | 1E4 |
| **ADAP Chromatogram Builder** | |
| a. Min group size in # of scans | 5 |
| b. Group intensity threshold | 5E4 |
| c. Min highest intensity | 5E4 |
| d. m/z tolerance – m/z | 0.005 |
| e. m/z tolerance - ppm | 0 |
| **Chromatogram Deconvolution** | |
| a. Algorithm | Wavelets (ADAP) |
| b. S/N Threshold | 200 |
| c. S/N estimator | Intensity window SN |
| d. Min feature height | 50000 |
| e. Coefficient/area threshold | 100 |
| f. Peak duration range | 0.10-1.0 |
| g. RT wavelet range | 0.01-0.20 |
| **RT Filtering** | |
| a. <60 seconds filtered out | |
| **Isotope Grouping** | |
| a. m/z tolerance – m/z | 0.005 |
| b. m/z tolerance – ppm | 0 |
| c. retention time tolerance | 0.5 min |
| d. maximum charge | 1 |
| e. representative isotope | Most intense |
| **Alignment** | |
| a. Method | Join aligner |
| b. m/z tolerance – m/z | 0.005 |
| c. m/z tolerance – ppm | 0 |
| d. weight for m/z | 75 |
| e. retention time tolerance | 0.5 min |
| f. weight for RT | 75 |
| **Blank Subtraction** | |
| a. Blank Files | 12 |
| b. Minimum # of detection | 5 |
| c. Quantification | Area |
| d. Ratio Type | Average |
| e. Fold change increase | 300% |
| **Gap filling** | |
| a. Method | Peak finder |
| b. Intensity tolerance | 10% |
| c. m/z tolerance – m/z | 0.005 |
| d. m/z tolerance – ppm | 0 |
| e. retention time tolerance | 0.25 |
